# Changes in endosymbiont complexity drive host-level compensatory adaptations in cicadas

**DOI:** 10.1101/353573

**Authors:** Matthew A. Campbell, Piotr Łukasik, Mariah M. Meyer, Mark Buckner, Chris Simon, Claudio Veloso, Anna Michalik, John P. McCutcheon

## Abstract

For insects that depend on one or more bacterial endosymbionts for survival, it is critical that these bacteria are faithfully transmitted between insect generations. Cicadas harbor two essential bacterial endosymbionts, *Sulcia muelleri* and *Hodgkinia cicadicola*. In some cicada species, *Hodgkinia* has fragmented into multiple distinct cellular and genomic lineages that can differ in abundance by more than two orders of magnitude. This complexity presents a potential problem for the host cicada, because low-abundance-but-essential *Hodgkinia* lineages risk being lost during the symbiont transmission bottleneck from mother to egg. Here we show that all cicada eggs seem to receive the full complement of *Hodgkinia* lineages, and that in cicadas with more complex *Hodgkinia* this outcome is achieved by increasing the number of *Hodgkinia* cells transmitted by up to six-fold. We further show that cicada species with varying *Hodgkinia* complexity do not visibly alter their transmission mechanism at the resolution of cell biological structures. Together these data suggest that a major cicada adaptation to changes in endosymbiont complexity is an increase in the number of *Hodgkinia* cells transmitted to each egg. We hypothesize that the requirement to increase the symbiont titer is one of the costs associated with *Hodgkinia* fragmentation.

## Introduction

Many organisms associate with microbial symbionts, in interactions that range from transiently pathogenic to stably beneficial from the host perspective. Beneficial symbionts can influence host biology in a variety of ways, but they often confer protection from natural enemies or provide nutrients to their hosts (1–5). Sap-feeding insects harbor obligate endosymbionts that supplement essential nutrients needed for normal host development and reproduction (1,6–9). For example, cicadas feed exclusively on nutritionally poor plant xylem sap (10,11), and therefore require supplementation with essential amino acids and vitamins (12). In many of the cicada species characterized to date (but see (13)), these nutritional services are provided by two transovarially transmitted bacterial endosymbionts, *Candidatus* Sulcia muelleri (hereafter referred to as *Sulcia*) and *Candidatus* Hodgkinia cicadicola (hereafter *Hodgkinia*) (14–16). We have previously shown that in two cicada genera, *Tettigades* and *Magicicada, Hodgkinia* has undergone an unusual form of lineage splitting (17–20). In some of these cicada species, a single *Hodgkinia* lineage has split into two or more derived lineages, each containing only a subset of the genes present in the single ancestral lineage. These reduced *Hodgkinia* genomes exist in separate cells and are in many cases complementary and partially non-redundant: each genome contains unique genes, and thus all are required to produce the same nutrients as the ancestral unsplit genome. The number of *Hodgkinia* lineages varies in different cicada species. For example, a species in the cicada genus *Diceroprocta* has one *Hodgkinia* lineage (21), various species in the genus Tettigades have between one and six *Hodgkinia* lineages (17,20), and the seven species in the long-lived periodical genus *Magicicada* contain more, possibly dozens, of *Hodgkinia* lineages (18,19).

A critical aspect of many symbiotic relationships is the transmission of symbionts between host generations. Mechanisms for symbiont transmission vary. Some organisms acquire symbionts from the environment each generation (22–24), while others have evolved mechanisms to transmit their symbionts directly to their offspring (9,25–30). We previously speculated that increases in *Hodgkinia* complexity might present intergenerational transmission problems for cicadas (18). As the number of *Hodgkinia* lineages increases, these lineages can start to vary in abundance by more than 100-fold in a single cicada (20). Hosts therefore risk losing the least abundant *Hodgkinia* lineages–which would likely result in inviable offspring–if they do not carefully manage the number and distribution of symbiont cells transmitted to each egg. We have hypothesized that cicadas with more complex *Hodgkinia* populations might compensate by increasing the number of *Hodgkinia* cells transmitted to each egg as a workaround to this problem (18). By contrast, we would not expect to see the same pattern for *Hodgkinia*’s partner symbiont, *Sulcia*, which has not been reported to increase in complexity. Finally, little is known about the mechanism of endosymbiont transfer in cicadas outside of work from the early 1900s, and nothing is known about how changes in *Hodgkinia* complexity may affect this process. Here we combine modeling, amplicon sequencing, and microscopy across cicada species and populations to study how increasing endosymbiont complexity affects symbiont transmission in cicadas.

## Methods

### Egg simulation protocol

For each of 1 to 30 hypothetical *Hodgkinia* cell lineages, between 1 and 2000 *Hodgkinia* cells were sampled with replacement in increments of 20 and placed in hypothetical eggs. If all lineages were present in the sample in at least one copy, that egg was determined to be viable. This procedure was repeated for all combinations of lineages and cell numbers, and the total proportion of viable eggs was calculated after 10,000 iterations. For the *T. chilensis* and *M. tredecim* experiments shown in Fig. 1B, the same simulation was performed but with the requirement that a minimum number of cells (1, 50, or 100) of each lineage be present in each egg for it to be deemed viable, as described in the results.

### Sample collection

Details of samples used for the study are shown in Table S1. For both *Tettigades* and *Magicicada* samples, all eggs in an “egg nest” were assumed to be laid by the same female. For *Tettigades* samples, we assumed that different nests were laid by different females because different egg nests were laid on different branches and the cicada population density was high where the samples were collected. In the case of *Magicicada*, we assumed that a series of adjacent egg nests on a single branch were produced by the same female. We attempted to verify this during data analysis, and as a precaution have removed any nests where eggs contained a different set of *Hodgkinia* genotypes than eggs in other nests in a series under the assumption that these may have been laid by a different female.

### DNA extraction

DNA from *M. septendecim* eggs and adult tissue, as well as Tettigades adult tissue, was extracted using a DNeasy Blood and Tissue kit (Qiagen, cat. #69506). DNA extraction process from *Tettigades* eggs was done by lysing the eggs in DNeasy lysis buffer followed by purification using Sera-Mag SpeedBeads (Carboxylate-Modified Particles, Thermo Scientific cat. # 09-981-123).

### Amplicon library preparation

Amplicon sequencing libraries were prepared following a two-step PCR protocol described in detail previously (20). For the first PCR step, we used primers targeting a gene retained on all (*Tettigades* spp. – *rpoB* with primers TCGCTRAGYTTAAYAAACGGATG and ATCGDTATTGCGMRGAGCTT) or some (*Magicicada* – *etfD* with primers ACGTTATTGTGGCYGAAGGTGC and ACGTTATTGTGGCYGAAGGTGC) *Hodgkinia* genomic circles present in a cicada, complete with Illumina adapters. During the second, indexing PCR step, additional adapters and sample-specific barcodes were added. The libraries were roughly quantified by comparison of band brightness following gel electrophoresis, pooled, and sequenced across three MiSeq lanes, alongside other libraries not included here. Sequencing for *Tettigades* was done across several MiSeq runs at the University of Montana Genomics Core, Missoula, MT. Sequencing for *Magicicada* was done on a MiSeq at the Genetic Resources Core Facility, Johns Hopkins Institute of Genetic Medicine, Baltimore, MD.

### Amplicon data analysis

The amplicon data were processed using mothur v. 1.39.5 (31). All reads were assembled into contigs, primer sequences were trimmed, and those reads with primer mismatches, ambiguous bases, homopolymer stretches >10 bp, or departing from the expected contig length by more than 10 bases were discarded. We then identified unique genotypes in the resulting filtered dataset, producing a table with information on the number of reads representing each genotype in each library. For the two *Tettigades* species, the exact sequences of *Hodgkinia* variants, alongside information on the relationship among and sequence diversity within cellular lineages, were available from our prior work (20). After verifying that no other abundant non-chimeric sequences were present within the table, we used only the counts of these exact genotypes for statistical comparisons. In the case of *M. septendecim*, we identified all genotypes that made up at least 1% of at least one library. The manual alignment and inspection of the sequences revealed that they represented two 99% OTUs that were about 7% divergent from each other. After manually identifying and discarding chimeric sequences among these OTUs, we used the count data for the remaining 37 genotypes, which together made up 83.0% of reads in a library on average (range 71.8-86.0%), for visualization and analyses.

Statistical comparisons of the lineage abundance among samples were conducted using R version 3.1.3 (32). Principal components analysis was conducted based on Bray-Curtiss dissimilarity matrices (functions vegdist and pco from packages vegan and labdsv, respectively) (33,34), and the results visualized using ggplot function (35). The multivariate analysis of variance among egg nests was conducted using the function adonis (package vegan (33)). The relative abundances of the two universally prevalent *Hodgkinia* genotypes among *Magicicada* egg nests were conducted using Generalized Linear Modeling, assuming quasibinomial error structure to account for overdispersion in the data.

### Microscopy

Fluorescent in-situ hybridization microscopy using small subunit rRNA probes was conducted on eggs as described previously (17). Briefly, eggs were broken manually, fixed for one hour in Carnoy’s solution, then incubated in prehybridization solution (12.5% dextran sulfate, 2.5X SCC, 0.25% BSA) at 37°C for 1 hr.Eggs were then briefly washed with warm 2XSCC and incubated overnight at 37°C with hybridization solution (prehybridization solution, 10ng/uL probe, 1.5ug/uL Hoechst 33258) in a humidity chamber. Eggs were then incubated in 2XSCC at 37°C for 1 hr, briefly rinsed with deionized H_2_O, placed on a glass slide, and covered with a cover slip. Probes used were Cy3-CCAATGTGGGGGWACGC for *Sulcia*, Cy5-CCAATGTGGCTGACCGT for *Hodgkinia* in *D. semicincta*, Cy5-CCAATGTGGCTGRCCGT for *Hodgkinia* in *Tettigades*, and Cy5-CCAATGTGGCTGTYCRT for *Hodgkinia* in *M. septendecim*. Symbiont balls in eggs were imaged on a Zeiss 880 confocal microscope. The total volume of the ball was estimated either as a sphere or spheroid. The number of *Sulcia* cells was counted within a box of approximately 50 × 50 × 10 micrometers^3^ within the tissue, and this number was used to estimate the total number of *Sulcia* cells present in the egg. The ratio of *Hodgkinia* to *Sulcia* cells present was then calculated on a single slice, and this value was used to estimate the number of *Hodgkinia* cells present. This process was repeated three times for each sample, and then averaged between samples.

For light microscopy, partially dissected cicada tissues were fixed in the field and stored in 0.05M phosphate-buffered solution with 2.5% glutaraldehyde, then fully dissected and postfixed using 1% osmium tetroxide, and embedded in Epon 812 (Serva, Germany) epoxy resin. Semi-thin sections (1 *μ*m thick) were stained with 1% methylene blue in 1% borax and analyzed and photographed under light microscope Nikon Eclipse 80i.

## Results

### Simulating the change to Hodgkinia cell transmission numbers

We first wanted to explore how changes in *Hodgkinia* complexity might affect the number of *Hodgkinia* cells transmitted from mother to egg from a theoretical perspective. Using computer simulations, we modeled transmission by first assuming that *Hodgkinia* lineages are transmitted from mother to egg randomly and that only a single cell of each *Hodgkinia* type is required for egg survival. Figure 1A shows the results for hypothetical cicadas harboring between one and thirty *Hodgkinia* lineages, with relative abundances based on the relative coverage values of completed genomic circles in the *M. tredecim* assembly (19). We find that as the *Hodgkinia* population becomes more complex, and especially as relative lineage abundances becomes more uneven, the minimum number of cells required so that all eggs are guaranteed to receive all *Hodgkinia* lineages grows quickly, by more than 2000-fold. We suspect that a 2000-fold increase is likely an upper bound on the changes we might expect to see, since we assume here that cicada eggs are viable if they only transmit one cell of any given lineage to each egg. Nevertheless, these results suggest that we could see up to orders-of-magnitude changes in *Hodgkinia* cell number transmission across a diversity of cicadas hosting *Hodgkinia* communities of varying complexities.

**Figure 1:**
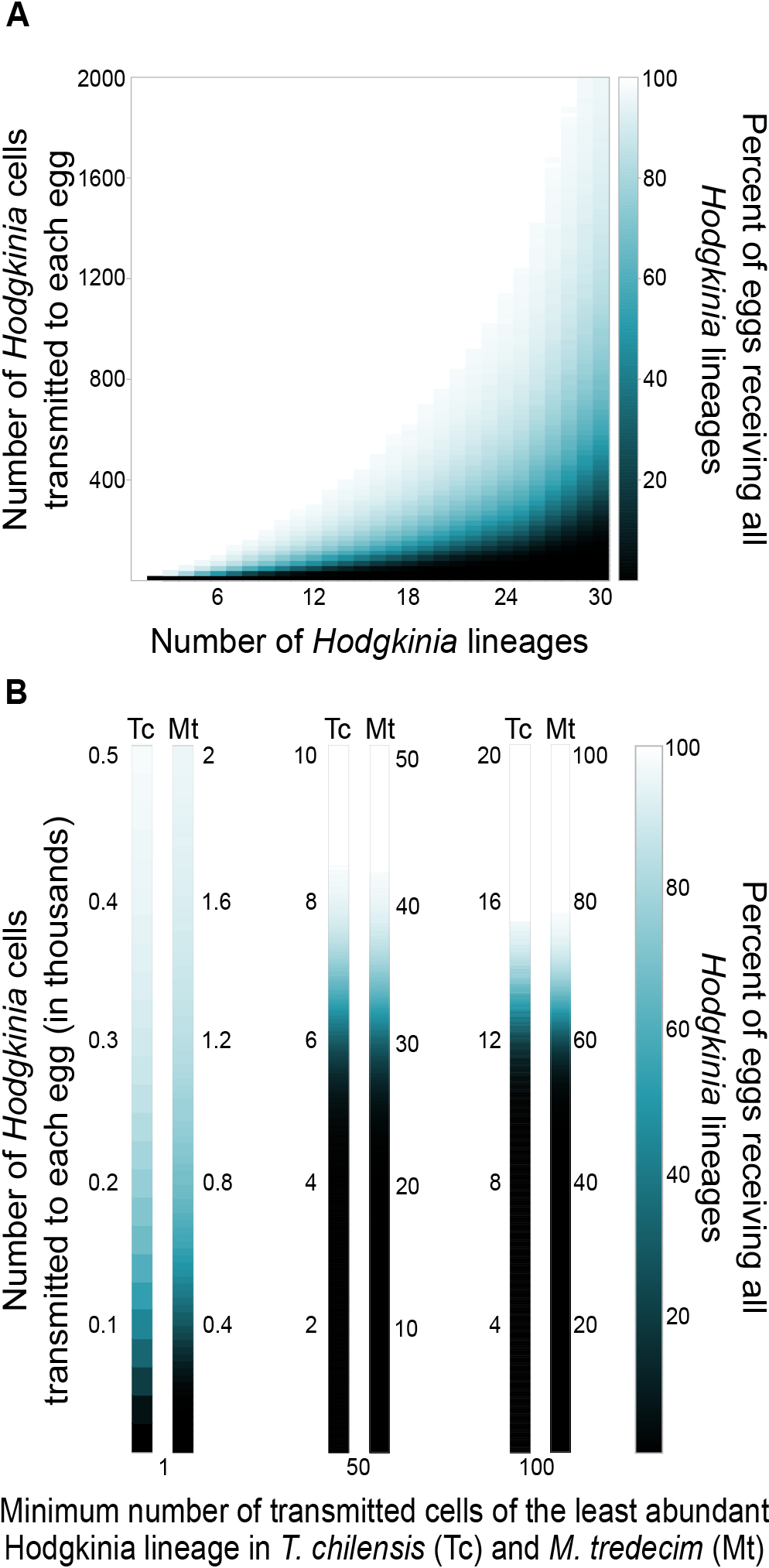
Simulation of the number of *Hodgkinia* cells required to be transmitted with increasing *Hodgkinia* complexity. A) Proportions of eggs receiving all *Hodgkinia* lineages for a given number of cells transmitted. Values for the abundance of the lineages were taken from sequencing coverages of the finished genomic circles in *M. tredecim* in (19). B) The same simulation for the six cellular lineages in *T. chilensis* (left-most bars) and many lineages in *M. tredecim* (right-most bars), requiring one (left), 50 (middle), or 100 (right) cells of the least abundant cellular lineage to be present in all eggs.

To get a sense of how the minimum number of *Hodgkinia* cells required for each lineage might affect changes in transmission number, we next modeled transmission in cicadas where we required a minimum of 1 single cell of each lineage in all eggs (Fig. 1B, left), 50 cells of each *Hodgkinia* lineage (Fig. 1B, middle), and 100 cells of each *Hodgkinia* lineage (Fig. 1B, right). These simulations used the *Hodgkinia* complexity of *T. chilensis* (6 lineages with a 69-fold abundance range) as well as *M. tredecim* (30 putative lineages with a 74-fold abundance range). For *T. chilensis*, requiring a single cell of each *Hodgkinia* lineage would necessitate that more than 500 *Hodgkinia* cells were transmitted to each egg. Requiring 50 cells of each *Hodgkinia* lineage would require that more than 8,000 cells are transmitted to each egg, and requiring 100 cells of each lineage would require over 15,000 *Hodgkinia* cells be transmitted to each egg. In each case for a cicada resembling *M. tredecim*, the host would need to transmit between 4- and 5-fold more *Hodgkinia* cells than in *T. chilensis*. These results suggest that we might see approximately five times more *Hodgkinia* cells transmitted in *M. tredecim* than *T. chilensis*.

**Figure 2:**
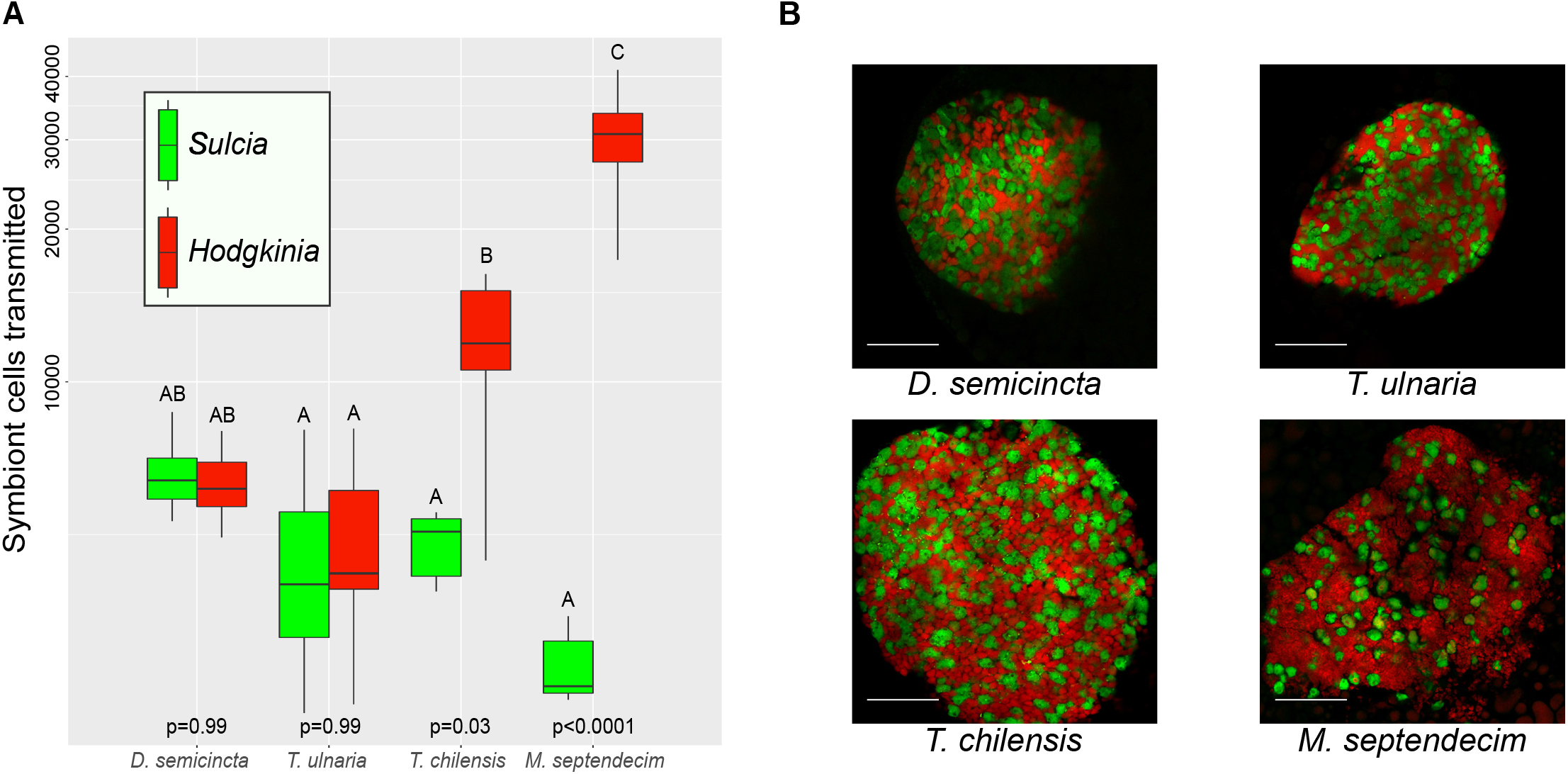
Numbers of symbiontcells transmitted to eggs in different cicadas. A) Boxplot of the number of *Sulcia* (green) and *Hodgkinia* (red) cells transmitted to eggs in *D. semicincta* (one lineage), *T. ulnaria* (one lineage), *T. chilensis* (six lineages), and *M. septendecim* (many lineages). Y axis uses a logarithmic scale. Reported p-values correspond to the test of whether more *Hodgkinia* than *Sulcia* cells are transmitted within a single species. B) Example images of the symbionts inside the eggs for the same four cicada species. Scale bars represent 50 microns.

### Cicadas harboring complex *Hodgkinia* populations transmit more Hodgkinia cells to eggs, but not more *Sulcia* cells

Our simulations show that the number of *Hodgkinia* cells transmitted to eggs is likely to increase with increasing *Hodgkinia* complexity. We tested this prediction by estimating the number of *Hodgkinia* cells transmitted to recently laid eggs from various cicada species (Fig. 2). We studied two distantly related cicada species with a single *Hodgkinia* lineage (*D. semicincta* and *T. ulnaria*), a species with six *Hodgkinia* lineages (*T. chilensis*), and a species with perhaps dozens of *Hodgkinia* lineages (*M. septendecim*). Using fluorescence microscopy, we first counted all of the *Hodgkinia* and *Sulcia* cells from a single confocal image slice. We then counted the number of *Sulcia* cells in a box of known volume and, modeling the symbiont ball as either a perfect sphere or spheroid, estimated the number of *Sulcia* cells in the entire symbiont ball. We then used the counted ratio of *Sulcia*:*Hodgkinia* to estimate the number of *Hodgkinia* cells present in the entire symbiont ball in the egg. We find that the average number of *Sulcia* cells transmitted to each egg varies approximately two-fold across all species, ranging from 2,572 in *M. septendecim* to 5,643 in *D. semicincta*, but that this difference is not statistically significant (Fig. 2A). In contrast, the numbers of *Hodgkinia* cells transmitted vary by as much as six-fold in different species, from 4,889 in *T. ulnaria* to 30,154 in *M. septendecim* (Fig. 2A). Within a cicada, the number of *Hodgkinia* cells differs significantly from *Sulcia* in *T. chilensis* (Tukey’s HSD p = 0.03) and *M. septendecim* (p < 0.0001), but not in *D. semicincta* or *T. ulnaria*. The transmitted *Hodgkinia.Sulcia* cell number ratio varies from ~1:1 in the cicadas with a single *Hodgkinia* lineage, to 2.4:1 in the species with six lineages, to 11.2:1 in the species harboring among the most complex *Hodgkinia* population known (Fig. 2B).

We estimated the number of transmitted cells of the least abundant *Hodgkinia* lineage by combining these total *Hodgkinia* cell estimates with our simulation data. Our simulations show that for *T. chilensis* to transmit 50 cells of the least abundant lineage, it would need to transmit between 8,000 and 9,000 total *Hodgkinia* cells, while for it to transmit 100 cells of the least abundant lineage it would need to transmit close to 16,000 total cells. We find that *T. chilensis* transmits approximately 12,000 *Hodgkinia* cells on average, and so we would expect it to transmit between 50 and 100 cells of the least abundant lineage. Using the same logic for *M. septendecim* (and again assuming all finished circles from (19) exist in different cells), which transmits approximately 30,000 total *Hodgkinia* cells, we would expect fewer than 50 cells of the least abundant *Hodgkinia* lineage to be present in each *M. septendecim* egg.

**Figure 3:**
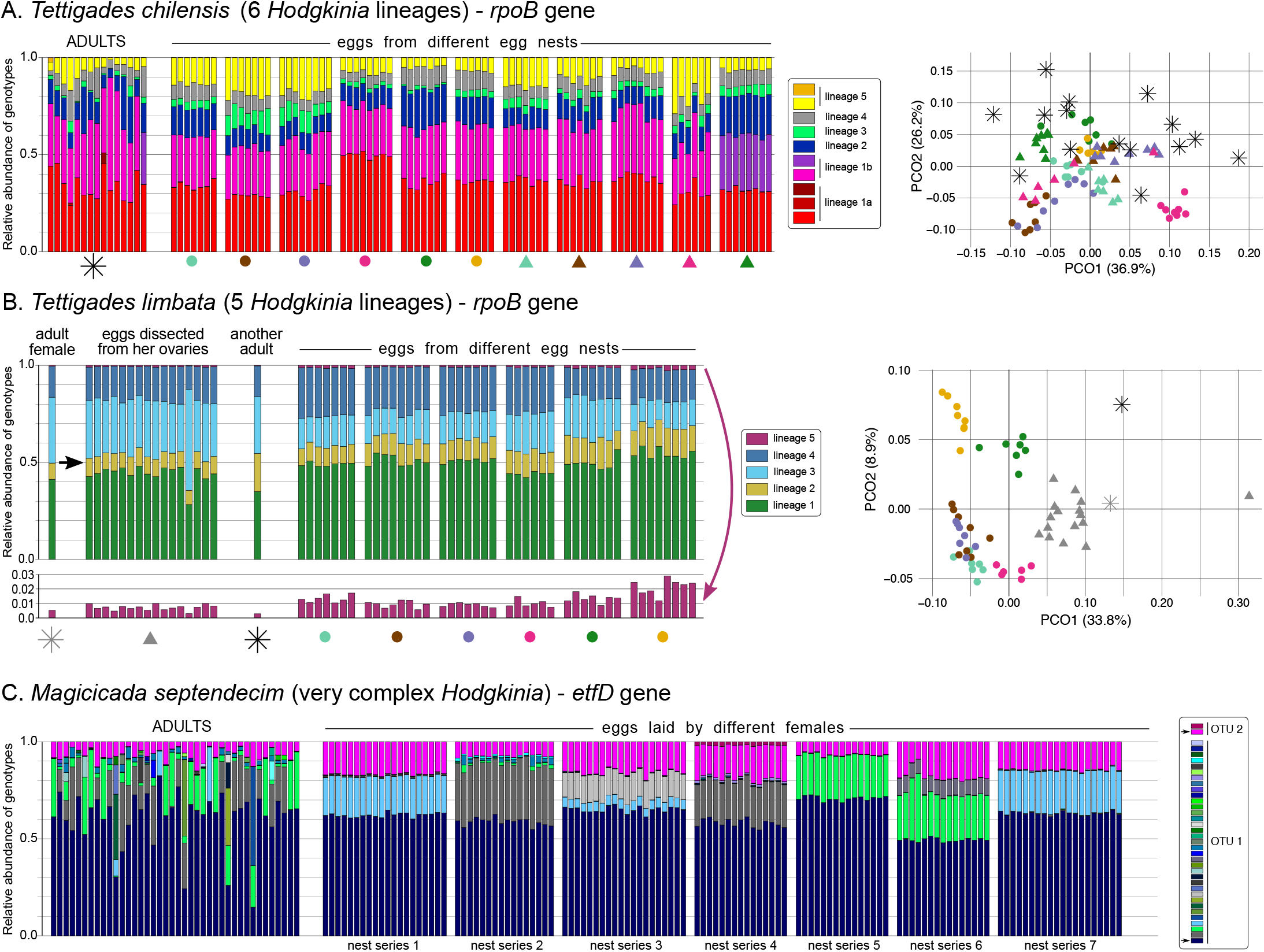
The relative abundances of *Hodgkinia* variants within populations of three cicada species, based on amplicon sequencing of symbiont-encoded protein-coding genes. For replicate adults and batches of eggs laid by individual females (egg nests), we plotted the relative abundance of *Hodgkinia rpoB* genotypes that correspond to six or five recognized lineages (*Tettigades* spp. - panels A & B); or of *Hodgkinia etfD* genotypes whose nature is less clear (*M. septendecim* - panel C). The relationships among samples of the two *Tettigades* species, based on the relative abundance of lineages rather than genotypes, is presented on Principle Components Analysis plots; shapes correspond to those shown below groups of barplots. In panel B, in a plot where scale is 10x magnified, we additionally show how the relative abundance of the rare lineage 5 varies among samples. In panel C, unique genotypes within the two observed OTUs are shown in shades of blue/green/grey (OTU1) or pink (OTU2), and those genotypes that are found in all samples are indicated with arrows on the legend.

### Cicada eggs seem to receive all *Hodgkinia* lineages, but variation in lineage abundances exists in the cicada population

Having shown that cicadas can adjust the number of symbiont cells transmitted to their eggs between species (Fig. 2), we next sought to measure how *Hodgkinia* lineages are transmitted between mother and eggs within and between species. We targeted protein-coding genes using amplicon sequencing to measure the differences in cell type abundances in eggs and in the bacteriome tissue of adult cicadas. For two *Tettigades* species, *T. chilensis* (6 cellular lineages) and *T. limbata* (5 cellular lineages), the target gene was RNA polymerase subunit B (*rpoB*), which is retained by all cellular lineages in all studied *Tettigades* species (20). Based on metagenomic data for single individuals (in the case of *T. chilensis*, from a divergent population), *rpoB* variants present in a cicada can vary by as much as 114-fold (20). In *Magicicada* species, gene targets were more difficult to choose because most assembled genomic circles encoded few genes and no single gene is universally conserved on each genome (19). We chose to target the electron transfer flavoprotein-ubiquinone oxidoreductase gene (*etfD*), which has two distinguishable gene homologs present at a 6-fold difference in abundance in *M. septendecim* (19).

We first assessed whether gene abundance estimates generated from amplicon sequencing were consistent between sequencing reactions and with genome abundance estimates we previously generated from metagenomics (19,20). We compared the abundance estimates for the two methods in three cicada species, and found that, in general, that the abundance estimates of genotypes obtained through amplicon sequencing were similar but not exactly the same as those found using metagenomics (Fig. S1A). In some cases, abundance estimates were very close (*T. chilensis*), while in others there was significant deviation in the relative abundance estimates for some lineages (*T. auropilosa* and *T. limbata*). Given that our genomic libraries were prepared using PCR-free methods or with <10 PCR cycles, and that our amplicon approach always required multiple (>25 in total) rounds of PCR with primers that might cause bias against some template variants, we assume that the proportions found using metagenomics are more accurate. Nevertheless, the abundance estimates found using amplicon data were consistent among technical replicates of the same sample (Fig. S1A) as well as between different parts of the bacteriome tissue from the same individual cicada (biological replicates – Fig. S1B), giving us confidence that the abundance differences we find between individuals result from genuine biological variation rather than methodological artifacts.

**Figure 4:**
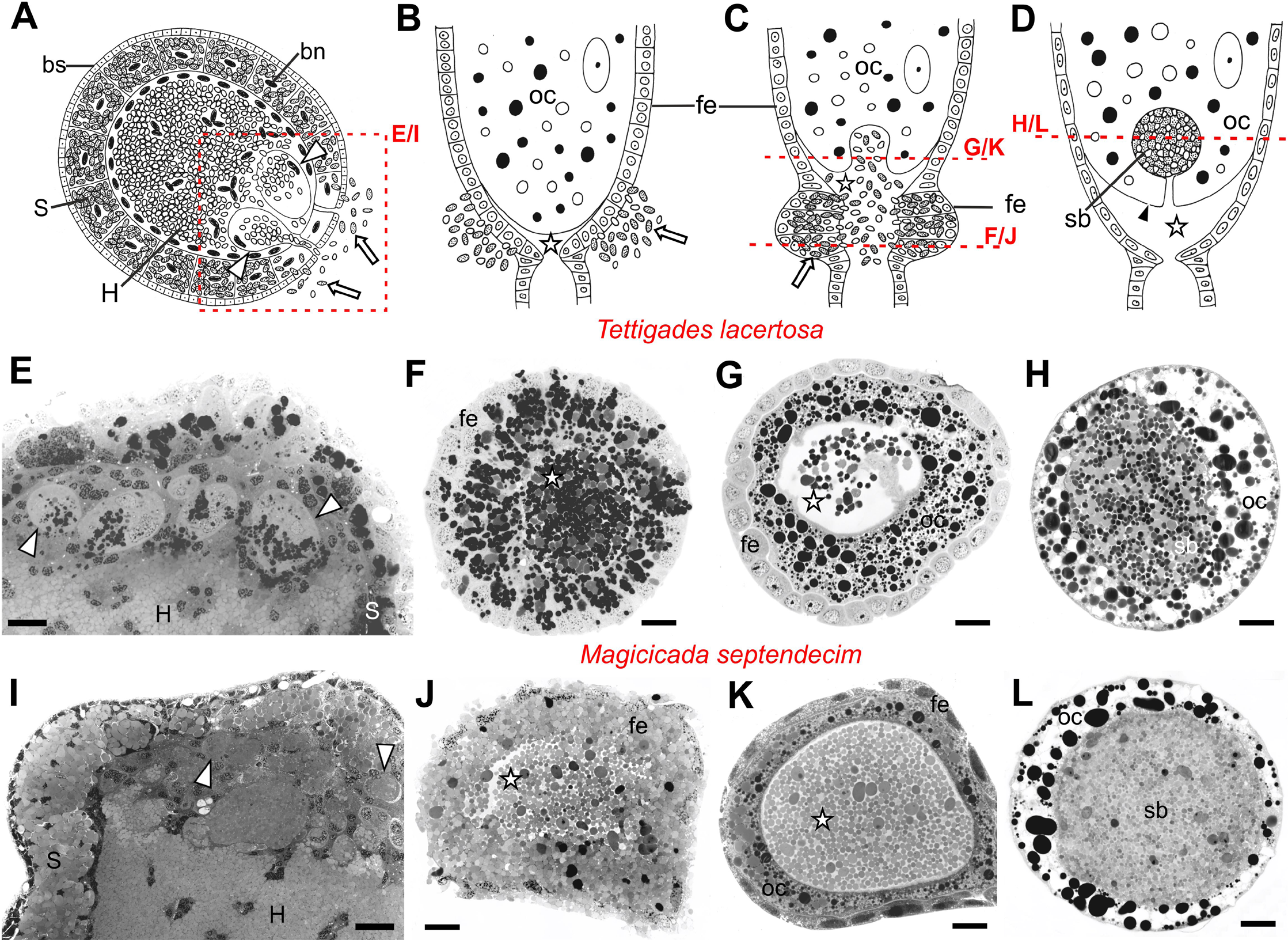
Transovarial transmission of endosymbiotic bacteria between cicada generations. A-D. Schematic representation of the successive stages of transmission, including the emigration of symbiont cells from the bacteriome (A), their migration through follicular epithelium into the perivitelline space of an ovariole (B-C), and then into an invagination within the basal part of the terminal oocyte (C) where they form a ‘symbiont ball’ (D). The microphotographs of methylene blue-stained sections indicated with red box or red line on the schematics are shown for two cicada species: *Tettigades lacertosa* which hosts three *Hodgkinia* lineages (E-H) and *Magicicada septendecim* that hosts very complex *Hodgkinia* (I-L). Note that the overall transmission process is the same in both species, but the numbers of migrating bacterial cells appear much greater in *Magicicada*. S – bacteriocyte with *Sulcia*; bn – bacteriocyte nucleus; bs – bacteriome sheath; H – syncytium with *Hodgkinia* cells; fe – follicular epithelium; oc – oocyte; sb – symbiont ball; white arrow – symbiotic bacterium; white arrowheads – *Hodgkinia*-carrying vesicles within syncytium; white star – perivitelline space; black arrowhead – oocyte membrane. Scale bar – 50 microns.

Our amplicon data revealed sequence complexity that was not detected in our previous metagenomic results (19,20). In *Tettigades limbata*, all specimens host the same *rpoB* genotypes that exactly correspond to sequences from our previous metagenomics work (20). The same is true in *T. chilensis*, except that in some cases one genotype has been replaced or complemented by another that differs by one nucleotide (Fig. 3A). In the case of *M. septendecim*, all sampled adults and eggs hosted two *Hodgkinia etfD* genotypes that were 6.7% divergent from each other at the nucleotide level (Fig. 3C). However, both amplicon sequences differed by one nucleotide substitution from the previously annotated *etfD* homologs in a metagenomic assembly of *M. septendecim* from a different brood (19). We suspect that these differences likely correspond to different alleles of the same *etfD* homologs. Additionally, all *M. septendecim* specimens hosted several genotypes that were less than 1% divergent from one of the two universally prevalent homologs (OTUs 1 and 2 in Fig. 3D). However, none of these derived genotypes are present in all samples, and all adults and egg nests harbor different combinations of derived genotypes.

We next tested whether cicadas reliably transmit all *Hodgkinia* lineages to each egg, and measured how the proportion of endosymbiont lineages varies within a single mother and within populations of single cicada species. Based on our simulation (Fig. 1) and cell count data (Fig. 2), we suspected that some cicada eggs might not receive all *Hodgkinia* lineages. Our amplicon data did not support this suspicion: we find that all *Tettigades* eggs contain all *rpoB* genotypes (Fig. 3A-B), and in *Magicicada*, all eggs contain both universally prevalent *etfD* genotypes (Fig. 3C). We then compared the variation in lineage proportions among adult cicadas, and batches of eggs laid by the females in the same populations. In Principal Components Analysis, *T. chilensis* eggs from the same nest tended to cluster together, separately from eggs from other nests, and the ADONIS test revealed significant differences in proportions of *Hodgkinia* lineages among eggs from the eleven characterized nests (F_10,68_ = 33.88, p < 0.001; Fig. 3A). In *T. limbata*, the differences in the proportions of lineages were less striking, but also significant among the six sampled egg nests (F_5,37_ = 30.16, p < 0.001; Fig. 3B). These differences were partly driven by the variable relative abundance of the least common lineage 5, which ranged among the studied samples over 10-fold (between 0.25% and 2.72%) (Fig. 3B).

We note that in *Magicicada* amplicons, a large number of unique genotypes complicates lineage abundance comparisons among samples. However, the comparisons of the relative abundance of the two universally prevalent *etfD* homologs revealed highly significant differences between egg batches from different females (GLM; genotype from OTU 1: F6,119=274.1, p < 0.001; genotype from OTU 2: F6,119=140.0, p < 0.001). We suspect that this sequence variation is the result of cicada population subdivision as well as some ancestral polymorphism in the cicada populations. There is some support for ancestral polymorphism in *Magicicada*: comparing the *etfD* genotype composition in individuals from different broods indicates that some of the variation is ancient and was present in the common ancestors of different broods (Fig. S2). Overall, the variation in lineage abundances that exists within cicada populations suggests that these insects can tolerate a relatively wide range of *Hodgkinia* lineage abundances. Individual mothers, however, seem to avoid substantial genotype abundance shifts between generations when transmitting symbionts to their offspring.

### The cell biological mechanism of symbiont transmission in cicadas is (mostly) conserved

Because we saw a clear adaptation by hosts in terms of changing the number of symbionts transferred in cicadas with varying levels of *Hodgkinia* complexity (Fig. 2), we wondered whether we could also observe changes to the mechanism of symbiont transfer. At the resolution of light microscopy, we find that the mechanism of endosymbiont transfer does not differ between *T. lacertosa* and *M. septendecim*, nor does it differ significantly from what Paul Buchner described in an unidentified African cicada species which appeared to harbor *Sulcia* and *Hodgkinia* (36) (Fig. 4). More generally, at this resolution, the mode of symbiont transmission appears well conserved throughout auchenorrhynchan insects (16,37). In mature cicada females, *Hodgkinia* and *Sulcia* cells are released from separate regions of the bacteriome into the hemolymph (Fig. 4A). Notably, *Hodgkinia* emigrates through large, nucleated subcellular compartments that form within the syncytium where it normally resides, while *Sulcia* is released directly from peripheral bacteriocytes. Subsequently, both bacterial symbionts migrate towards the ovarioles and through follicular cells into the perivitelline space (Fig. 4B-C). As the number of symbionts in that space increases, the oocyte membrane creates a deep invagination where the symbionts gather. Later, as the opening closes, the intermixed *Sulcia* and *Hodgkinia* cells form a characteristic ‘symbiont ball’ in each egg (Fig. 4D).

The transmission process does not appear to be qualitatively different between *Tettigades* (Fig. 4E-H) and *Magicicada* (Fig. 4I-L). However, consistent with our fluorescent microscopy observations (Fig. 2A), in *Magicicada* the overall number of bacterial cells migrating into the oocyte is visibly higher than in *Tettigades*, and the ratio of *Hodgkinia* cells to *Sulcia* cells is higher than in *Tettigades* (Fig. 2B). Together, these data indicate that in response to *Hodgkinia* splitting, cicadas have adjusted their ancient transmission pathway to increase the numbers of transmitted *Hodgkinia* cells, but not *Sulcia* cells.

## Discussion

### Cicadas adapt to increases in *Hodgkinia* complexity

The strong selective pressure to reliably transmit nutritional symbionts to offspring is reflected in a conserved mechanism for transmission in cicadas. In *D. semicincta* and *T. ulnaria*, cicada species diverged by tens of million years (38–41), both *Sulcia* and *Hodgkinia* have stable, conserved genomes (17,21), and we have shown here that these two cicadas also transmit similar numbers of *Hodgkinia* and *Sulcia* cells to each egg (Fig. 2A). Within the last ~4 million years, *Hodgkinia* in some *Tettigades* species has become more complex due to lineage splitting and genome reduction (17,20). This same process had led to the incredibly complex situation seen in all *Magicicada* species, which we estimate has been ongoing over the last 5-20 million years (19).

This increase in symbiont complexity poses a problem for the cicada. Rather than transmitting a single lineage each of *Sulcia* and *Hodgkinia*, the cicadas with more complex *Hodgkinia* must now transmit *Sulcia* plus many distinct–but still essential–*Hodgkinia* lineages. This problem has three obvious and not mutually exclusive solutions. Solution 1:The host evolves a mechanism to distinguish between *Hodgkinia* lineages and actively places all lineages into each egg. Because the *Hodgkinia* genome no longer encodes the machinery to make its own membranes, the host must define *Hodgkinia*’s envelope, so this solution is formally possible. Solution 2: The host could increase the number of *Hodgkinia* cells transmitted to each egg, thereby increasing the odds that lower abundance lineages make it to each egg. Solution 3: The host mother could produce some proportion of (presumably inviable) eggs that do not receive all *Hodgkinia* lineages. This last option would obviously come with a huge negative fitness cost for the host.

We currently do not have the ability to measure whether hosts actively select certain *Hodgkinia* lineages (solution 1). We do find that cicadas seem to be able to tolerate substantial variation in *Hodgkinia* lineage abundances (Fig. 3), suggesting that if a host selection process does happen then it is not highly accurate over cicada generations. We find clear evidence that hosts increase the number of *Hodgkinia* cells transmitted to eggs (solution 2, Fig. 2), but no evidence that any egg is missing any *Hodgkinia* lineages (solution 3, Fig. 3). From these data, we conclude that the increase in symbiont transmission number is likely a key adaptation by the cicada to compensate for *Hodgkinia*’s increasing complexity. The increase in *Hodgkinia* transmission numbers appears to solve this aspect of the symbiont complexity problem, since all cellular lineages seem to be reliably transmitted to all offspring (Fig. 3) We note however that it is possible that some low abundance lineages are occasionally lost in certain eggs and that we lack the sensitivity to see it.

Individual *Hodgkinia* lineages can differ in abundance by more than 100-fold in adult cicadas (20). Since eggs receive similar proportions of the lineages that were present in their mother (Fig. 4), the least abundant lineages will be the primary drivers of the required increase of transmitted *Hodgkinia* cells. Because it seems unlikely that cicadas can indefinitely increase the number of *Hodgkinia* cells transmitted to each egg, cicadas must also decrease the number of cells transmitted of the least abundant *Hodgkinia* lineage. Our simulations estimate that *T. chilensis* and *M. septendecim* might receive fewer than 100 cells of the least abundant *Hodgkinia* lineage (Fig. 1). These estimates are consistent with our expectation based on relative sequencing coverage: we estimate that *T. chilensis* eggs receive only ~84 cells of the least abundant lineage (based on sequencing coverage for *T. chilensis* of a different population, where its equivalent comprises 0.8% of the total *Hodgkinia* population (20)), and *M. septendecim* eggs likely receive fewer than 50 cells of the least abundant lineage.

The ability to decrease the amount of the least abundant lineage is only possible because cicadas with single *Hodgkinia* lineages transmit substantially more *Hodgkinia* cells than is strictly necessary (Fig. 2). This “surplus” of transmitted cells acts as a buffer for *Hodgkinia* lineage splitting, and this buffer is likely the reason we see only a ~6-fold increase in *Hodgkinia* cells transmitted as *Hodgkinia* complexity increases, rather than the ~2,000-fold increase seen in our simulations (Fig. 1A). The relatively smaller increase that we measure empirically (Fig. 2) vs. that which we predict computationally (Fig. 1) might also be due to more than one *Hodgkinia* genomic circle sharing cellular lineages (20). Our genomic data strongly suggest that at least in the genus *Tettigades*, some *Hodgkinia* genomic circles are present in the same *Hodgkinia* cell, but we have not yet verified this result using other methods (20). While reducing the minimum number of required cells is one method to prevent the required transmission size from spiraling out of control, we also know that lineage splitting in at least some cicadas is ongoing (19). Therefore, the lower cell number distribution limit is not something that cicadas can continue reducing indefinitely. For example, the cobalamin biosynthesis gene *cobQ* is only encoded by 0.8% of all *Hodgkinia* cells in *T. chilensis* (20), so further decrease in the abundance of the *cobQ*-bearing lineage may negatively affect the supply of this vitamin.

### *Hodgkinia* is driving the adaptation

Importantly, we have shown that the number of *Sulcia* cells transmitted remains relatively stable in all of the studied cicadas [and may even be lower in *Magicicada* (Fig. 2A), though the decrease is not statistically significant]. We thus infer that the principal driver of the transmission changes we show here is specific to *Hodgkinia*-related processes rather than a general change in host transmission strategy. It is also formally possible that *Hodgkinia*’s transmission numbers could have changed before *Hodgkinia* started splitting, and thus be the enabling the fragmentation we see in some cicadas. The transmission numbers for *Sulcia* and *Hodgkinia* in cicadas with unsplit *Hodgkinia* lineages are on the high end for transovarially transmitted symbionts estimated for a wide range of other Hemipteran insects (Table 1), but this alone seems unlikely to be the main driver of lineage splitting in *Hodgkinia* because some cicadas continue to retain *Hodgkinia* with a single genome structure.

Though the increase in *Hodgkinia* transmission number solves the cicadas’ immediate problem, it raises other potential complications. Cicadas, including *Magicicada*, typically lay between 400-600 eggs (48,49), but *M. septendecim* individuals transmit ~6-fold more *Hodgkinia* cells to each egg than *D. semicincta* or *T. ulnaria* individuals. If a cicada is to continue transmitting larger numbers of *Hodgkinia* cells to all eggs, it must either lay fewer eggs, continually replenish its *Hodgkinia* population as it lays eggs, or maintain a larger *Hodgkinia* population in its adult stage. Laying fewer eggs is likely to lead to fewer offspring so is unlikely to be favored. It may be possible for cicada mothers to replenish the *Hodgkinia* population as she lays eggs, because Buchner has suggested that *Hodgkinia* may be dividing prior to transmission into eggs (36). However, our microscopy shows no clear evidence of this (Fig. 4), so it is unclear if this is an important mechanism for increasing *Hodgkinia* numbers. This mechanism would also require relatively rapid *Hodgkinia* reproduction since cicadas lay their eggs within a short time span (50). While not definitive, we have also gathered anecdotal evidence that cicadas with more complex *Hodgkinia* populations harbor larger *Hodgkinia* populations as adults (18), but we currently have no solid data on the total number of symbiont cells in adult cicadas. But maintaining a larger *Hodgkinia* population would bring its own complications, as the cicada has to provide more tissue space and nutrients for a larger *Hodgkinia* population, and runs the risk of crowding out its partner symbiont *Sulcia* (Fig. 2, (18)).

**Table 1.**
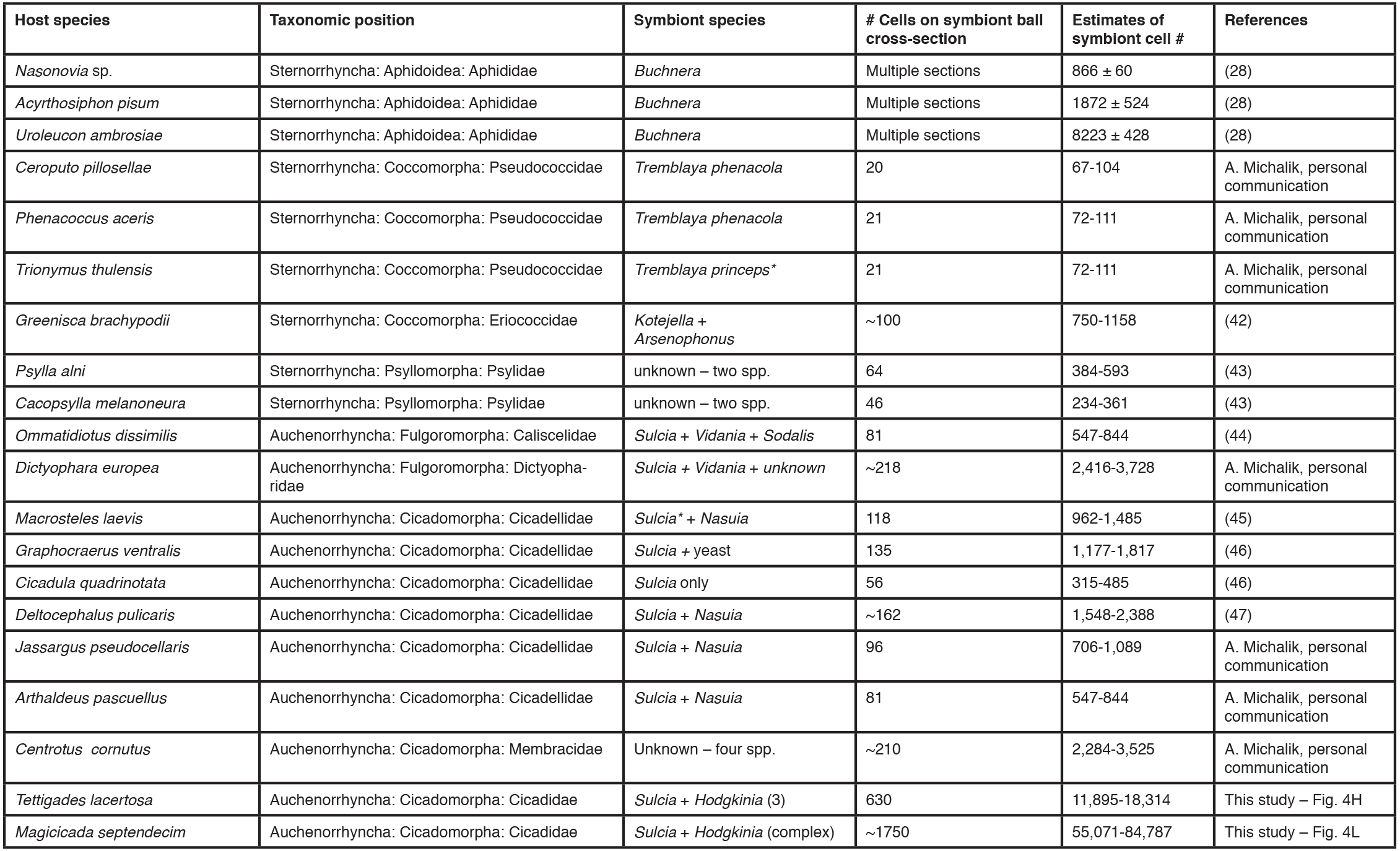
The estimated numbers of endosymbiont cells within symbiont balls in eggs of different hemipteran species. Because bacterial species are sometimes hard to distinguish, cells of different species were counted together. For species other than aphids, the number is based on cell count on a single cross section, with the assumption that the ball was spherical and symbionts evenly distributed within the ball. The lower estimate is based on assumption that the section was made through the centre of a spherical symbiont ball; the higher estimate assumes that the section was made at 25% of the ball length. Note that these estimates may be inaccurate if the section was made even closer to the ball edge, or if the shape of the ball departed significantly from spherical. *Cells of endosymbionts of two species contain endobacterial symbionts, which were not included in the counts.

### Symbiont population sizes could affect host- and symbiont-levels of selection

An increase in *Hodgkinia*’s intra-cicada population size may have implications for the long-term evolution of the symbiosis. As in any endosymbiosis, the evolutionary trajectories of host and symbiont are not inevitably and permanently aligned. In order for the host to better control the evolution of its symbiont, it is important that hosts maintain their symbionts at small effective population sizes, which is often achieved by subjecting symbionts to strong population bottlenecks at transmission (51–54). Maintaining small intra-host symbiont effective population sizes does three things. First, it reduces the efficacy of symbiont-level selection for selfish traits, since selection is less efficacious in small populations. Second, small symbiont populations will harbor less diversity, further decreasing the efficacy of symbiont-level selection. Finally, with relatively few symbionts within a cicada, there are fewer mutational targets to acquire the complementary gene loss required for *Hodgkinia* splitting to happen. While speculative, it seems possible that increasing the number of *Hodgkinia* cells transmitted could make the splitting process more likely to happen, because it would decrease the level of control that the host can impart on its symbionts. Larger symbiont populations would lead to more intra-host variation, and thus more chances for lineage splitting by mutation and drift or by symbiont-level cheating as previously hypothesized (17–19). In this scenario, the host increasing the load of *Hodgkinia* cells might lead to a positive-feedback loop, where the compensatory changes cicadas have evolved to deal with degenerative *Hodgkinia* evolution might make the problem of splitting worse.

It is perhaps unsurprising that symbiont evolution is driving compensatory adaptations in cicadas. There are a number of other examples of what appears to be host compensatory evolution to symbiont change, such as nuclear genes responding to high mitochondrial substitution rates in the plant genus *Silene* (55,56) and primates (57), horizontal transfer of bacterial genes to the nucleus to maintain symbiont function in several eukaryotic groups (reviewed in (58)), and the evolution of trafficking systemsto move gene products between host and symbiont (59–61). These examples highlight the pervasiveness of hosts compensating for the evolution of symbiont traits, and might reflect the peril of hosts critically relying on vertically transmitted endosymbionts (62–64): if endosymbionts erode in functionality due to host restriction and genetic drift, the host must compensate somehow or suffer the consequences of reduced fitness or, in extreme cases, extinction.

## Acknowledgements

We thank DeAnna Bublitz for collecting the D. semicincta eggs used in this study, and all members of the McCutcheon lab for helpful discussion. We also thank Art Woods for suggesting the egg simulation heatmap, Lou Herritt and the Molecular Histology and Fluorescence Imaging Core at the University of Montana for help with imaging the eggs, and Ada Jankowska for help with drawing the scheme in Figure 4. This study was supported by National Science Foundation grants IOS-1256680, IOS-1553529, and DEB-1655891, by the National Aeronautics and Space Administration Astrobiology Institute Award NNA15BB04A, by the National Geographic Society grant 9760-15, and by the American Genetics Association EECG Research Award 2015. CS acknowledges additional support from the University of Connecticut.

## Data availability

The amplicon sequencing data have been deposited in GenBank, under BioProject accessions PRJNA475285, PRJNA475287, and PRJNA476567. Accession numbers for individual libraries are provided in Table S2.

## References

1. Moran NA, Baumann P. Bacterial endosymbionts in animals. Curr Opin Microbiol 2000; 3(3):270–5.

2. Moran NA, McCutcheon JP, Nakabachi A. Genomics and Evolution of Heritable Bacterial Symbionts. Annu Rev Genet 2008; 42(1):165–90.

3. Douglas AE. Mycetocyte symbiosis in insects. Biol Rev Camb Philos Soc. 1989; 64(4):409–34.

4. Wernegreen JJ. Genome evolution in bacterial endosymbionts of insects. Nat Rev Genet 2002; 3(11):850–61.

5. Douglas AE. How multi-partner endosymbioses function. Nat Rev Microbiol 2016; 14(12):731–43.

6. Wang J-J, Dong P, Xiao L-S, Dou W. Effects of removal of Cardinium infection on fitness of the stored-product pest Liposcelis bostrychophila (Psocoptera: Liposcelididae). J Econ Entomol 2008; 101(5):1711–7.

7. Kuriwada T, Hosokawa T, Kumano N, Shiromoto K, Haraguchi D, Fukatsu T. Biological role of Nardonella endosymbiont in its weevil host. PLoS One 2010; 5(10):e13101.

8. Ishikawa H, Yamaji M. Symbionin, an aphid endosymbiont-specific protein–I. Insect Biochemistry 1985; 15(2):155–63.

9. Hosokawa T, Hironaka M, Inadomi K, Mukai H, Nikoh N, Fukatsu T. Diverse strategies for vertical symbiont transmission among subsocial stinkbugs. PLoS One 2013; 8(5):e65081.

10. White J, Strehl CE. Xylem feeding by periodical cicada nymphs on tree roots. Ecol Entomol 1978; 3(4):323–7.

11. Lloyd M, White J. Xylem Feeding by Periodical Cicada Nymphs on Pine and Grass Roots, With Novel Suggestions for Pest Control in Conifer Plantations and Orchards. Ohio J Sci 1987; 87(3):50–4.

12. Christensen H, Fogel ML. Feeding ecology and evidence for amino acid synthesis in the periodical cicada (Magicicada). J Insect Physiol 2011; 57(1):211–9.

13. Matsuura Y, Moriyama M, Łukasik P, Vanderpool D, Tanahashi M, Meng X-Y, et al. Recurrent symbiont recruitment from fungal parasites in cicadas. Proc Natl Acad Sci U S A 2018; 47:201803245.

14. McCutcheon JP, McDonald BR, Moran NA. Convergent evolution of metabolic roles in bacterial co-symbionts of insects. Proc Natl Acad Sci U S A 2009; 106(36):15394–9.

15. Moran NA, Tran P, Gerardo NM. Symbiosis and insect diversification: an ancient symbiont of sap-feeding insects from the bacterial phylum Bacteroidetes. Appl Environ Microbiol 2005; 71(12):8802–10.

16. Buchner P. Endosymbiosis of Animals with Plant Microorganisms. Wiley: New York, USA, 1965.

17. Van Leuven JT, Meister RC, Simon C, McCutcheon JP. Sympatric Speciation in a Bacterial Endosymbiont Results in Two Genomes with the Functionality of One. Cell 2014; 158(6):1270–80.

18. Campbell MA, Van Leuven JT, Meister RC, Carey KM, Simon C, McCutcheon JP. Genome expansion via lineage splitting and genome reduction in the cicada endosymbiont Hodgkinia. Proc Natl Acad Sci U S A 2015; 112(33):10192–9.

19. Campbell MA, Łukasik P, Simon C, McCutcheon JP. Idiosyncratic Genome Degradation in a Bacterial Endosymbiont of Periodical Cicadas. Curr Biol 2017; 27(22):3568–3575.e3.

20. Łukasik P, Nazario K, Van Leuven JT, Campbell MA, Meyer M, Michalik A, et al. Multiple origins of interdependent endosymbiotic complexes in a genus of cicadas. Proc Natl Acad Sci U S A 2018; 115(2):E226–35.

21. McCutcheon JP, McDonald BR, Moran NA. Origin of an Alternative Genetic Code in the Extremely Small and GC-Rich Genome of a Bacterial Symbiont. PLoS Genet 2009; 5(7):e1000565.

22. Nussbaumer AD, Fisher CR, Bright M. Horizontal endosymbiont transmission in hydrothermal vent tubeworms. Nature 2006; 441(7091):345–8.

23. Kikuchi Y, Hosokawa T, Fukatsu T. Insect-microbe mutualism without vertical transmission: a stinkbug acquires a beneficial gut symbiont from the environment every generation. Appl Environ Microbiol 2007; 73(13):4308–16.

24. Nyholm SV, McFall-Ngai MJ. The winnowing: establishing the squid-vibrio symbiosis. Nat Rev Microbiol 2004; 2(8):632–42.

25. Ereskovsky AV, Bouryesnault N. Cleavage pattern in Oscarella species (Porifera, Demospongiae, Homoscleromorpha): transmission of maternal cells and symbiotic bacteria. J Nat Hist 2002; 36(15):1761–75.

26. Al-Khalifa MS. The transovarial transmission of symbionts in the grain weevil, Sitophilus granarius. J Invertebr Pathol 1984; 44(1):106–8.

27. Eberle MW, McLean DL. Observation of symbiote migration in human body lice with scanning and transmission electron microscopy. Can J Microbiol 1983; 29(7):755–62.

28. Mira A, Moran NA. Estimating Population Size and Transmission Bottlenecks in Maternally Transmitted Endosymbiotic Bacteria. Microb Ecol 2002; 44(2):137–43.

29. Koga R, Meng X-Y, Tsuchida T, Fukatsu T. Cellular mechanism for selective vertical transmission of an obligate insect symbiont at the bacteriocyte-embryo interface. Proc Natl Acad Sci U S A 2012; 109(20):E1230–7.

30. Hosokawa T, Kikuchi Y, Fukatsu T. How many symbionts are provided by mothers, acquired by offspring, and needed for successful vertical transmission in an obligate insect-bacterium mutualism? Mol Ecol 2007; 16(24):5316–25.

31. Schloss PD, Westcott SL, Ryabin T, Hall JR, Hartmann M, Hollister EB, et al. Introducing mothur: Open-Source, Platform-Independent, Community-Supported Software for Describing and Comparing Microbial Communities. Appl Environ Microbiol 2009; 75(23):7537–41.

32. Team RC. R: A language and environment for statistical computing. Vienna, Austria; 2013. Available from: http://R-project.org/

33. Oksanen J, Kindt R, Legendre P, OHara B. The vegan package. Community ecology package [Internet]. 10:631–7. Available from: http://cran.r-project.org/web/packages/vegan/index.html

34. Roberts DW. labdsv: Ordination and Multivariate Analysis for Ecology. 1:3–1. Available from: http://CRAN.R-project.org/package=labdsv

35. Wickham H. ggplot2: Elegant Graphics for Data Analysis [Internet]. Springer-Verlag New York; 2016. Available from: http://ggplot2.org

36. Buchner P. Studien an intracellularen Symbionten. I: Die intracellularen Symbionten der Hemipteren. Arch. Protistenk 1912; 26:1–116.

37. Szklarzewicz T, Michalik A. Transovarial Transmission of Symbionts in Insects. In: Oocytes. Cham: Springer International Publishing; 2017. pp. 43–67.

38. Cooper KW. Davispia bearcreekensis Cooper, a new cicada from the Paleocene, with a brief review of the fossil Cicadidae. Am J Sci 1941; 239(4):286–304.

39. Kritsky G, Poinar G Jr. Morphological conservatism in the foreleg structure of cicada hatchlings, Novicicada burmanica n. gen.,n. sp. in Burmese amber, N. youngin. gen., n. sp. in Dominican amber and the extant Magicicada septendecim (Fisher) (Hemiptera: Cicadidae). Hist Biol 2012; 24(5):461–6.

40. Poinar G Jr, Kritsky G, Brown A. Minyscapheus dominicanus n. gen., n. sp. (Hemiptera: Cicadidae), a fossil cicada in Dominican amber. Hist Biol 2012; 103:1–5.

41. Marshall DC, Hill KBR, Moulds M, Vanderpool D, Cooley JR, Mohagan AB, et al. Inflation of Molecular Clock Rates and Dates: Molecular Phylogenetics, Biogeography, and Diversification of a Global Cicada Radiation from Australasia (Hemiptera: Cicadidae: Cicadettini). Syst Biol 2016; 65(1):16–34.

42. Michalik A, Szwedo J, Stroiński A, Świerczewski D, Szklarzewicz T. Symbiotic cornucopia of the monophagous planthopper Ommatidiotus dissimilis (Fallén, 1806) (Hemiptera: Fulgoromorpha: Caliscelidae). Protoplasma 2018; 50:1–13.

43. Kot M. Struktura i rozwój jajnika oraz transowarialny przekaz endosymbiotycznych mikroorganizmów u koliszków (Insecta, Hemiptera: Psylloidea). [Kraków]. 2015.

44. Michalik A, Schulz F, Michalik K, Wascher F, Horn M, Szklarzewicz T. Coexistence of novel gammaproteobacterial and Arsenophonus symbionts in the scale insect Greenisca brachypodii (Hemiptera, Coccomorpha: Eriococcidae). Environ Microbiol 2018; 20(3):1148–57.

45. Kobiałka M, Michalik A, Walczak M, Junkiert Ł, Szklarzewicz T. Sulcia symbiont of the leafhopper Macrosteles laevis (Ribaut, 1927) (Insecta, Hemiptera, Cicadellidae: Deltocephalinae) harbors Arsenophonus bacteria. Protoplasma. 2015; 253(3):903–12.

46. Kobiałka M, Michalik A, Walczak M, Szklarzewicz T. Dual “Bacterial-Fungal” Symbiosis in Deltocephalinae Leafhoppers (Insecta, Hemiptera, Cicadomorpha: Cicadellidae). Microb Ecol 2017; 75(3):771–82.

47. Kobiałka M, Michalik A, Walczak M, Junkiert Ł, Szklarzewicz T. Symbiotic microorganisms of the leafhopper Deltocephalus pulicaris (Fallén, 1806) (Insecta, Hemiptera, Cicadellidae: Deltocephalinae): molecular characterization, ultrastructure and transovarial transmission. Pol J Entomol 2015; 84(4):143–304.

48. Itô Y, Nagamine M. Why a cicada, Mogannia minuta Matsumura, became a pest of sugarcane: an hypothesis based on the theory of “escape.” Ecol Entomol 1981; 6(3):273–83.

49. Karban R. Effects of local density on fecundity and mating speed for periodical cicadas. Oecologia 1981; 51(2):260–4.

50. Williams KS, Simon C. The ecology, behavior, and evolution of periodical cicadas. Annu Rev Entomol 1995; (40):269–95.

51. O’Fallon B. Population structure, levels of selection, and the evolution of intracellular symbionts. Evolution 2008; 62(2):361–73.

52. Rispe C, Moran NA. Accumulation of Deleterious Mutations in Endosymbionts: Muller’s Ratchet with Two Levels of Selection. Am Nat 2000; 156(4):425–41.

53. Pettersson ME, Berg OG. Muller’s ratchet in symbiont populations. Genetica 2006; 130(2):199–211.

54. Bastiaans E, Aanen DK, Debets AJM, Hoekstra RF, Lestrade B, Maas MFPM. Regular bottlenecks and restrictions to somatic fusion prevent the accumulation of mitochondrial defects in Neurospora. Philos Trans R Soc Lond, B 2014; 369(1646):20130448–8.

55. Sloan DB, Triant DA, Wu M, Taylor DR. Cytonuclear interactions and relaxed selection accelerate sequence evolution in organelle ribosomes. Mol Biol Evol 2014; 31(3):673–82.

56. Sloan DB. Using plants to elucidate the mechanisms of cytonuclear co-evolution. New Phytol 2015; 205(3):1040–6.

57. Osada N, Akashi H. Mitochondrial-nuclear interactions and accelerated compensatory evolution: evidence from the primate cytochrome C oxidase complex. Mol Biol Evol 2012; 29(1):337–46.

58. Husnik F, McCutcheon JP. Functional horizontal gene transfer from bacteria to eukaryotes. Nat Rev Microbiol 2018; 16(2):67–79.

59. Nowack ECM, Grossman AR. Trafficking of protein into the recently established photosynthetic organelles of Paulinella chromatophora. Proc Natl Acad Sci U S A 2012; 109(14):5340–5.

60. Nakabachi A, Ishida K, Hongoh Y, Ohkuma M, Miyagishima S-Y Aphid gene of bacterial origin encodes a protein transported to an obligate endosymbiont. Curr Biol 2014; 24(14):R640–1.

61. Singer A, Poschmann G, Mühlich C, Valadez-Cano C, Hänsch S, Hüren V, et al. Massive Protein Import into the Early-Evolutionary-Stage Photosynthetic Organelle of the Amoeba Paulinella chromatophora. Curr Biol 2017; 27(18):2763–5.

62. Bennett GM, Moran NA. Heritable symbiosis: The advantages and perils of an evolutionary rabbit hole. Proc Natl Acad Sci U S A 2015; 112(33):10169–76.

63. Keeling PJ, McCutcheon JP. Endosymbiosis: The feeling is not mutual. J Theor Biol 2017; 434:75–9.

64. Kiers ET, West SA. Evolution: Welcome to Symbiont Prison. Curr Biol 2016; 26(2):R66–8.

